# Device Design and Diagnostic Imaging of Radiopaque 3D Printed Tissue Engineering Scaffolds

**DOI:** 10.1101/2024.02.26.582070

**Authors:** Mitchell Delemeester, Kendell M. Pawelec, Jeremy M.L. Hix, James R. Siegenthaler, Micah Lissy, Philippe Douek, Angèle Houmeau, Salim Si-Mohamed, Erik M. Shapiro

## Abstract

3D printed biomaterial implants are revolutionizing personalized medicine for tissue repair, especially in orthopedics. In this study, a radiopaque Bi_2_O_3_ doped polycaprolactone (**PCL**) composite is developed and implemented to enable the use of diagnostic X-ray technologies, especially photon counting X-ray computed tomography (**PCCT**), for comprehensive in vivo device monitoring. PCL filament with homogeneous Bi_2_O_3_ nanoparticle (**NP**) dispersion (0.8 to 11.7 wt%) are first fabricated. Tissue engineered scaffolds (**TES**) are then 3D printed with the composite filament, optimizing printing parameters for small feature size and severely overhung geometries. These composite TES are characterized via micro-computed tomography (*µ***CT**), tensile testing, and a cytocompatibility study, with Bi_2_O_3_ mass fractions as low as 2 wt% providing excellent radiographic distinguishability, improved tensile properties, and equivalent cytocompatibility of neat PCL. The excellent radiographic distinguishability is validated in situ by imaging 4 and 7 wt% TES in a mouse model with µCT, showing excellent agreement with in vitro measurements. Subsequently, CT image-derived swine menisci are 3D printed with composite filament and re-implanted in their corresponding swine legs *ex vivo*. Re-imaging the swine legs via clinical CT allows facile identification of device location and alignment. Finally, the emergent technology of PCCT unambiguously distinguishes implanted menisci in situ.

## 1 Introduction

The use of implantable biomedical tissue engineering scaffolds (**TES**) in the clinic is growing rapidly [1]. Often, TES are engineered to perform critical functions in regenerative medicine, including providing mechanical support, delivering therapeutic molecules, and creating protective barriers. Clinically approved TES for these purposes include devices such as hernia patches and orthopedic devices [45, 46]. As TES gain clinical acceptance, the trend is to adapt these tools for personalized medicine. This has been demonstrated most successfully for orthopedic devices that can be 3D printed on demand and often at the point-of-care, to repair bone structures after trauma [10]. In preclinical studies of 3D printed personalized TES, fused filament fabrication (**FFF**) is frequently employed for device production as it is highly accessible and low cost [10, 11]. Of the materials that can be 3D printed, poly(caprolactone) (**PCL**) is biocompatible, has excellent toughness, and has an *in vivo* degradation rate of 3-4 years [38, 39], thus making it a promising material for on demand TES. However, while PCL is compatible with FFF printing, limited printing parameters are available in literature which do not adequately take model geometry into account [15, 40].

Despite clinical success, the use of TES poses inherent risks to patients, including misalignment and premature degradation associated with infection and inflammation [41]. The ability to monitor devices post-implantation would mitigate these risks by enabling rapid post-surgical realignment or later post-surgical intervention. Post-implantation however, TES manufactured from organic polymers cannot be visually inspected or palpated and are poorly distinguished from native tissue with available clinical imaging modalities. In a simulated clinical trial, we reported that radiologists have ≈ 80% success rate in identifying polymer TES implant location but *<* 55% success rate in assessing damage, regardless of imaging technique (magnetic resonance imaging (**MRI**), computed tomography (**CT**), ultrasound) [2]. In the case of orthopedic device failure, the inability to monitor implants can lead to delays in treatment that reduce patient mobility and impact quality of life [28].

To monitor TES performance, radiologists need to: 1) identify implant location, 2) accurately discriminate it from the surrounding tissue, and 3) evaluate implant integrity. The embedding of biocompatible radiopaque nanoparticles (**NP**s) in polymer TES has been demonstrated to effectively impart radiopacity, rendering TES easily detectable by x-ray computed tomography (**CT**) in pre-clinical models [3, 4]. TES fabricated from these radiopaque nanocomposites significantly increased radiologist success when monitoring their integrity to *>* 85% [2]. Radiopaque nanoparticles have been utilized successfully in the past with compositions of noble metals like gold and platinum [36], and various metal oxides [4, 10, 16]. Bismuth oxide, which has low cost and a long history of use in the medical field [37], is very well suited for CT because of its high Z number (83) and high linear X-ray attenuation coefficient [9, 20].

CT is a ubiquitous imaging technology in the clinic with high spatial resolution and rapid acquisition time relative to MRI [36], making it an ideal choice for monitoring TES. Further, the emergent technology of photon counting CT (**PCCT**) enables the discrimination of high atomic number elements (Z*>*40) from surrounding tissue in pre-clinical settings [5]. PCCT translation to the clinical setting is expected to reduce radiation dose and improve image quality, enabling enhanced imaging techniques involving multiple material analysis [6]. Bismuth, which is not naturally found in the body, has been previously identified as an excellent candidate for use with PCCT [7]. While coated Bi_2_O_3_ NP contrast agents have already been utilized in PCCT studies [8], the use of uncoated Bi_2_O_3_ NPs in additively manufactured PCL devices to monitor *in vivo* device performance has not been previously investigated.

In this study we merged the concepts of personalized TES with advanced clinical monitoring, implementing PCCT to push the boundaries of what is currently possible. To accomplish this goal, Bi_2_O_3_ NPs were compounded with PCL and extruded as filament for FFF, enabling the fabrication of highly radiopaque TES. 3D printed scaffolds were highly biocompatible and clinically sized devices exhibited excellent radiopacity with contrast to noise ratios that allowed them to be readily distinguished from surrounding tissues in situ. Further, we demonstrated that the implants could be easily segmented from bone in orthopedic applications via the emergent technology of PCCT.

## 2 Results and Discussion

### 2.1 Homogeneous Radiopaque Filaments

TES that enable *in situ* monitoring require homogeneous radiopacity throughout the structure, ensuring that any detectable changes observed in the structure are the result of physical changes to the device. To create radiopaque filament for 3D printing, Bi_2_O_3_ NPs (216 ± 37 nm Fig. 1 b-c) were combined with Facilan™ PCL 100 filament at nominal concentrations of 0.5-15 wt% (Fig. 1 a). Inductively coupled plasma mass spectrometry (ICP-MS) revealed the actual Bi_2_O_3_ composition of the fabricated filaments was 0.2 to 11.7 wt%. This slight deviation from nominal percentages has been observed in other studies as well and is likely due to the mixing process [12].

**Figure 1:**
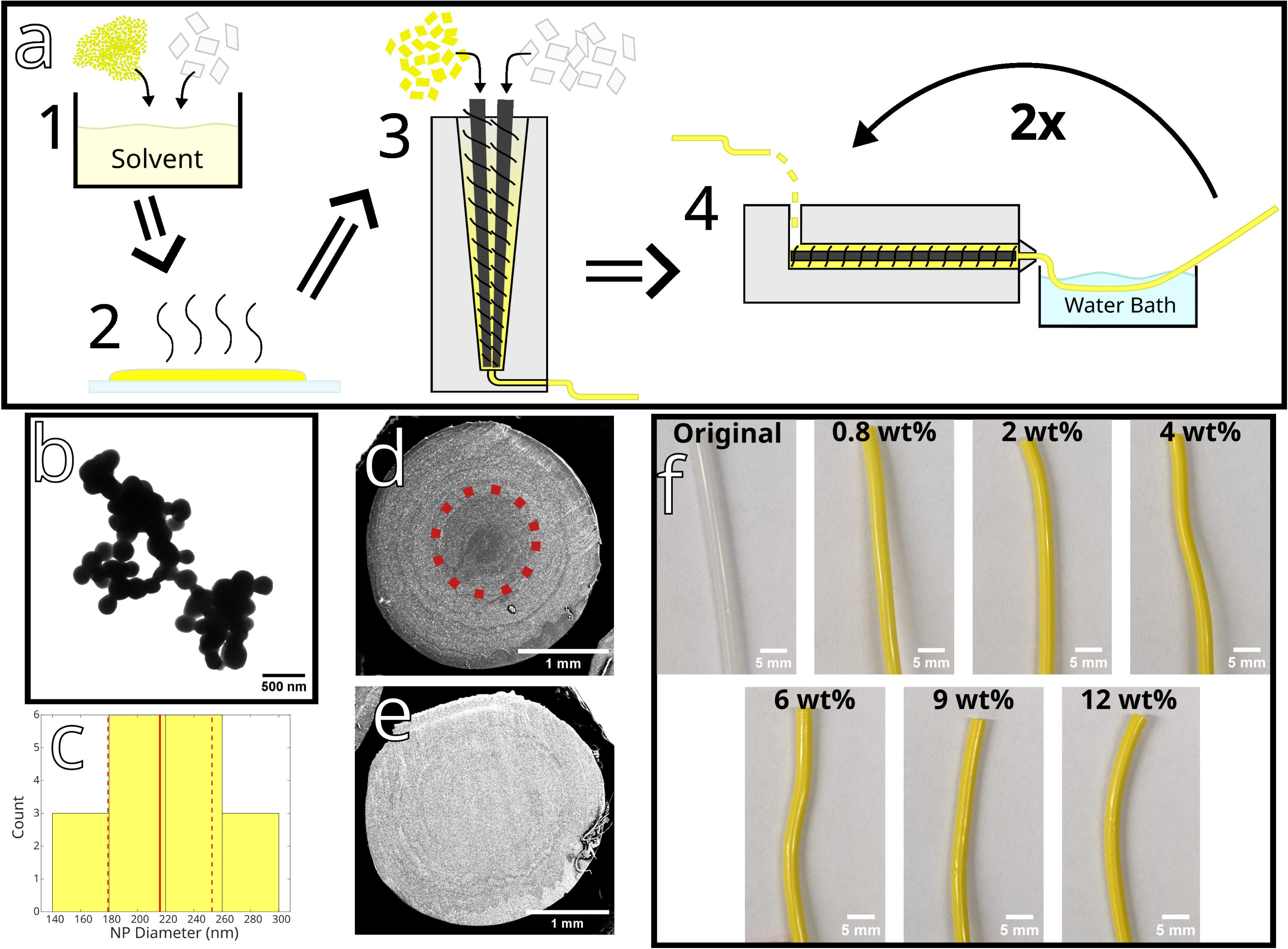
Homogeneous, radiopaque filaments were produced from composites of PCL and Bi_2_O_3_ NPs. Schematic of the process **(a)** to make dimensionally accurate filament with homogenous NP dispersion. NPs imaged with transmission electron microscopy (TEM) were spherical **(b)** with an average size of 216 ± 37 nm **(c)**. Filament was checked for homogeneity via scanning electron microscopy (SEM) backscatter electron micrograph. The cross section of an inhomogeneous 4 wt% filament with a residual core (circled in red) is shown in **(d)**, compared to homogenous 7 wt% filament in **(e)**. Composite filaments **(f)** ranged from 0 wt% (white) to 12 wt% (yellow) Bi_2_O_3_.

The NP homogeneity in the fabricated filaments was verified with scanning electron microscopy backscatter imaging. As Bi_2_O_3_ NP concentration is linearly correlated with X-ray attenuation [12], producing homogeneous filament has a direct impact on the radiopacity of a TES and its monitoring post-implantation. NP distribution in filament must be verified post-production before being used in 3D printing. Initial runs of filament were found to have an inhomogeneous distribution of NPs, as seen in the cross-section of the filament in Fig. 1 d. When this shell & core structure was formed in the filament (Fig.1 a.4), it remained even after subsequent printing. To eliminate inhomogeneity in the batches, each batch was run through the final extrusion process (Fig. a.4) at least twice.

### 2.2 Optimization of 3D Printing TES with PCL

PCL is a biocompatible polymer with many desirable properties for medical devices [11]. While it is compatible for FFF printing, limited printing parameters are available in literature. Material properties such as melting temperature and heat transfer are important factors in printing TES at high resolution. Not only is PCL difficult to print, given its low melting temperature compared to other polyesters, but changes in melt characteristics with NP addition must also be considered. Initial 3D printing with PCL filament revealed that insufficient cooling of the deposited bead resulted in unstable features and print failures. To understand the melt and solidification characteristics of both the neat and composite filaments, differential scanning calorimetry (**DSC**) was employed. The crystallization temperature (*T*_c_) of the neat PCL (0 wt% Bi_2_O_3_) was found to be 29° C, very close to room temperature (Fig. 2a). Adding Bi_2_O_3_ NPs increased the *T*_c_ of all the composites, ranging from 33° C to 35° C as the concentration increased from 0.8 wt% to 11.7 wt%. In contrast, the melt temperature of the composites and neat PCL both aligned with the published literature value of 60 C [13].

**Figure 2:**
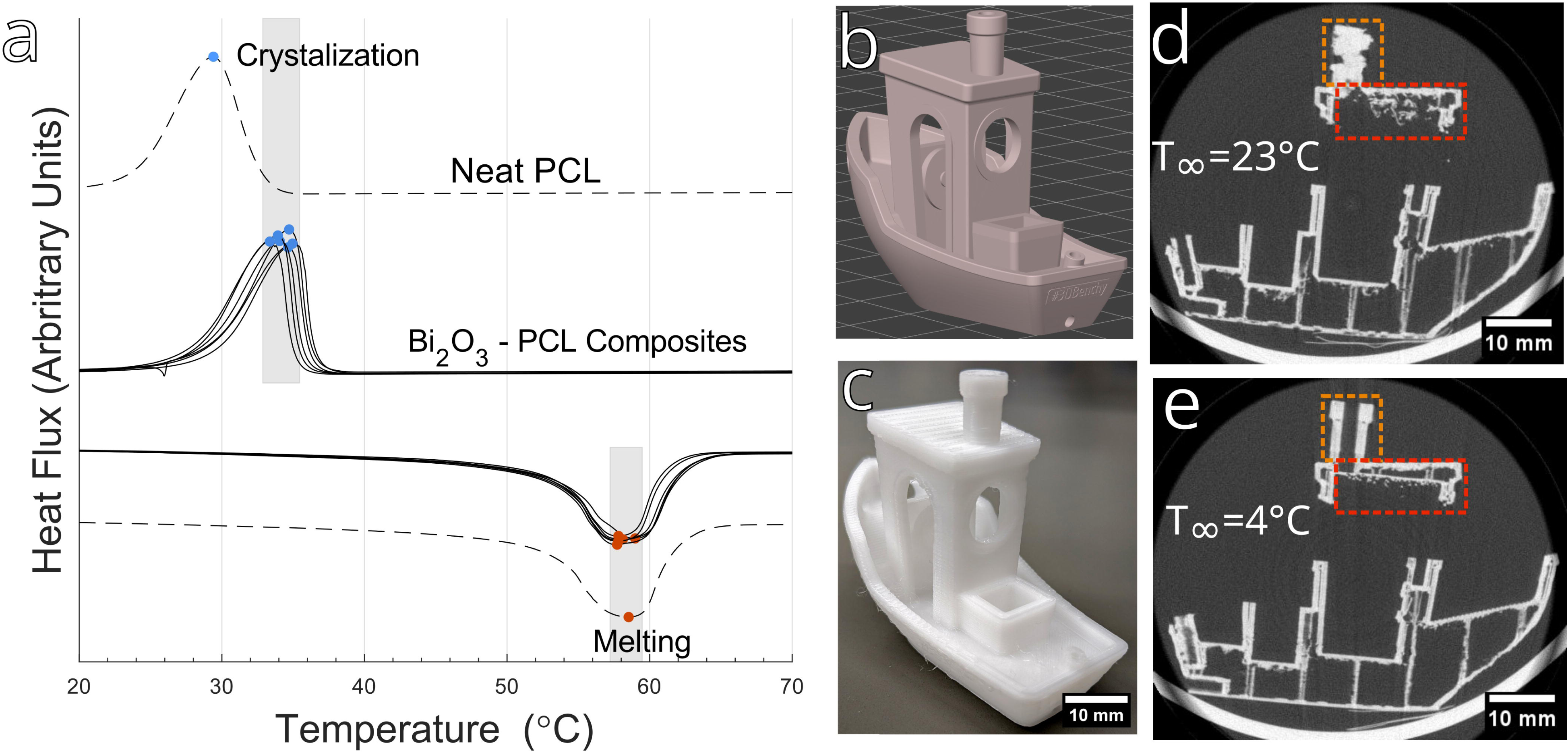
Optimization of PCL printing relied on controlling heat flux. **(a)** DSC curves of neat PCL (dashed) and composites (solid). **(b)** Benchy 3D model used for testing a broad range of 3D printing parameters. **(c)** 3D printed Benchy from neat PCL using the optimized settings and printed in a cold room. **(d)** µCT of a Benchy printed at room temperature ambient conditions. **(e)** µCT of the same print performed at 4°C and at ambient temperature. The boxes encompass features that failed at ambient room temperature and printed successfully in a cold room.

To address 3D printing in the worst-case scenario (lowest *T*_c_), neat PCL filament was selected for 3D printing optimization with a standard benchy 3D model [47], Fig. 2 b-e. During testing, two scenarios were identified as very difficult to 3D print due to the long solidification time of the bead: unsupported overhangs (<55° from horizontal), and layers with short print times (*<*60 s). Differences in printing parameters for these cases are summarized in Table 1. Nozzle diameter and temperature remained constant in all scenarios, while speed for print moves, ambient temperature, and use of supports were found to be most critical for success in other scenarios. Optimized speeds for print moves, or “carriage speed”, for specific nozzle temperature were taken from Ortega et al [40]. In addition, when printing in a cold room with a low ambient temperature (4° C), increasing the bed temperature was found to improve part adhesion to the print bed.

**Table 1.** Optimized Printing Parameters for FFF of PCL using a 0.3mm nozzle.

| Scenarios | Ambient Temp (°C) | Nozzle Dia. (mm) | Nozzle Temp (°C) | Speeds for Print Moves (mm s<sup>-1</sup>) | Fan Speed (%) | Bed Temp (°C) | Supports (Y/N) |
| --- | --- | --- | --- | --- | --- | --- | --- |
| Typical | 23 | 0.3 | 110 | 10 | 100 | 30 | N |
| Small Cross Sections | 23 | 0.3 | 110 | 3-9 | 100 | 30 | N |
| Unsupported Overhangs (<55°) | 23 | 0.3 | 110 | 10 | 100 | 30 | Y |
| Cold Room | 4 | 0.3 | 110 | 10 | 50-100 | 35 | N |

The close *T_c_* of neat PCL to ambient temperature resulted in failed prints with short layer times or unsupported overhangs. From classical heat transfer, it’s known that convection and radiation from a surface depend on a temperature differential to drive heat transfer [14]. With a sufficiently small bead, as in this case, the surface and internal temperature of the bead may be considered the same. The total heat transfer from convection and radiation can then be written as:

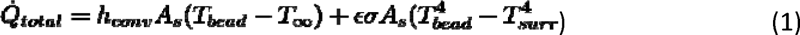

where *T_bead_* is the temperature of the deposited bead, *A_s_* is the area of the surface, *h* and IZ and *σ* are constants, and *T*_∞_ and *T_surr_* are determined by ambient conditions. As Δ*T* decreases, the convective heat transfer decreases linearly and the radiative heat transfer decreases as a 4th power. In room temperature ambient conditions, the bead therefore remains in a viscoelastic liquid state longer than necessary while the *T_c_* is slowly approached. By decreasing the ambient temperature around the printed bead, Δ*T* remains sufficiently large to quickly drive the bead below *T*_c_. This in turn improves printing performance of overhangs and short layer times.

To verify this experimentally, an identical object was printed with the same parameters in both room temperature (T=24° C) and cold room (T=4°C) environments. When printing in a cold room, significant reductions in print deformation and improved performance in bridging gaps were observed (Fig. 2, d-e). These results align with previous findings by Paetzold et al., who showed that 3D printing PCL in an ambient temperature of 17.5° C produced scaffolds with better geometrical fidelity than printing in an ambient temperature of 27.5° C [15].

However, 3D printing at 4°C ambient conditions posed other problems. Bed adhesion decreased with decreasing temperature, as the higher temperature differential increased thermal shrinkage, especially for larger parts (build plate footprint > 10 cm^2^ ). Additionally, the printer would enter an error state between prints due to cold circuitry, requiring frequent returns of the printer to room temperature ambient conditions. 3D printing in 4° C ambient conditions is therefore only suggested when printing small TES (build plate footprint < 10 cm^2^ ) and/or when using support material for overhangs is not feasible. Additional temperature ranges between 4 and 17.5° C should be tested to determine the optimal ambient temperature for 3D printing PCL.

### 2.3 Tensile Properties of Radiopaque TES

Having determined optimized parameters for printing PCL and PCL-NP composites, the range of Bi_2_O_3_ NPs that could be incorporated into PCL had to be defined. This was accomplished by characterizing 1) mechanical properties, 2) biocompatibility, and 3) x-ray attenuation. Understanding the effect of Bi_2_O_3_ NPs on composites aids in the selection of compositions for specific applications, as the mechanical properties of a TES greatly influence their performance *in vivo*.

Mechanical testing was employed to investigate the effect of Bi_2_O_3_ wt% on the tensile properties of as-printed TES. Reference tensile samples, fabricated to assess as-printed TES performance, are shown in Fig. 3 c (before) and d (after) testing. The tensile samples were designed to represent TES with 60% porosity. The tensile samples typically failed near the grips, although some samples failed near the center, Fig. 3 d. Statistically significant increases of the apparent yield stress and apparent elastic modulus were found for low wt% composites (≤ 2 wt%). The greatest increase in properties occurred at 0.8 wt% Bi_2_O_3_ (apparent yield stress = 961 ± 28 kPa, apparent elastic modulus = 1.86 ± 0.005 kPa) corresponding to increases of 32% and 55% over neat PCL, respectively (Fig. 3 a). No change in yield strain was found for any composites compared to neat PCL.

**Figure 3:**
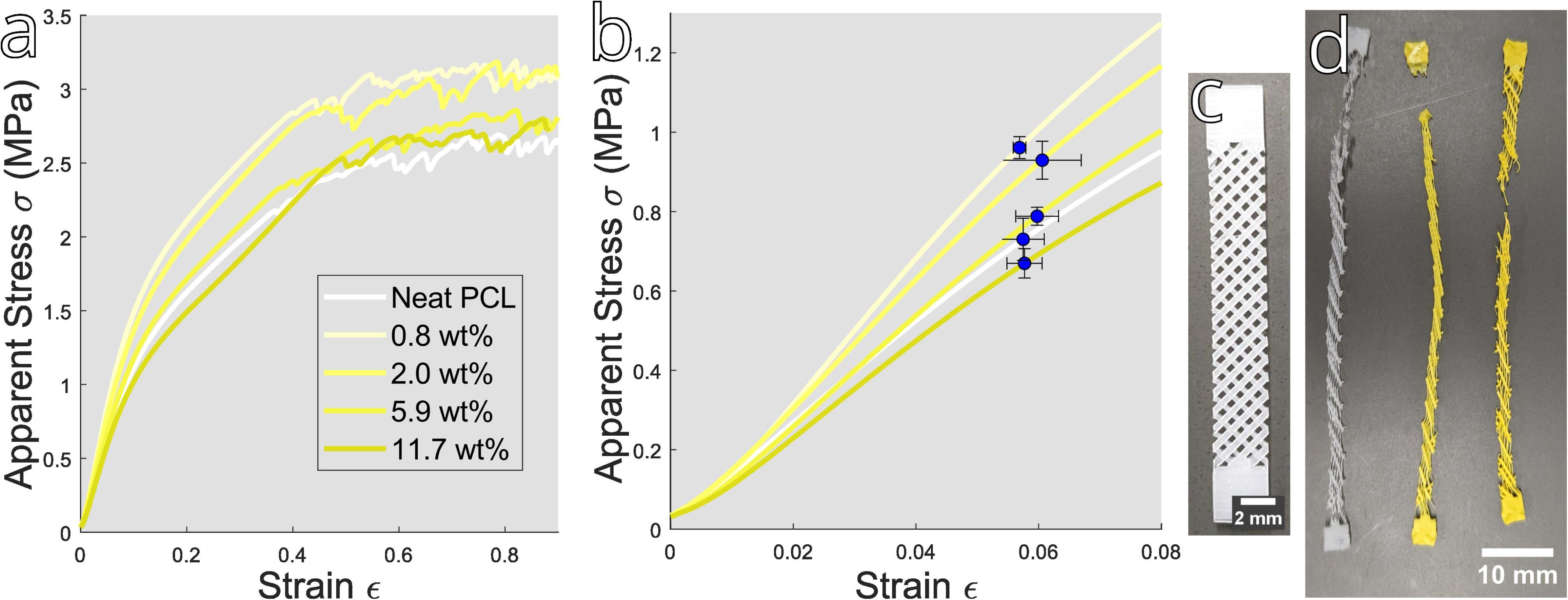
Tensile properties of PCL filaments are affected by the addition of radiopaque Bi_2_O_3_ nanoparticles and melt processing. Tensile properties of composites; a typical Stress-Strain curve **(a)** is plotted with the mean apparent yield stress and yield strain (± standard deviation) shown in **(b)**. Tensile testing was performed on specimens with 40% infill: **(c)** before testing and **(d)** after testing. Scale bar: (b) 2 mm, (c) 10 mm.

Typically, the elastic modulus and yield stress of a composite increase with NP content [32]. Interfacial interactions between the polymer matrix and NPs result in a large interphase volume. Previous research has shown that changes in elastic modulus are dominated by the properties of the interphase [32]. Thus, the increase in stiffness of the composites with < 2 wt% NPs agrees with the trends seen in literature. However, in other systems the elastic modulus continues to increase with much higher NP loadings. For example, barium sulfate NPs in poly(L-lactide) increased the elastic modulus by 62% at a loading of 15 wt%, before dropping off at 20 wt% [16]. In PCL composited with magnesium oxide NPs, the peak elastic modulus (+280%) was achieved at the maximum tested loading of 30 wt% [17]. In our case, the occurrence of maximum modulus and yield stress is at significantly lower NP loadings.

Other factors were investigated to understand the drop-off of composite elastic moduli at low NP loadings. The crystallinity of the composites as a function of NP loading was estimated from the heat of crystallization from DSC data (Fig. 2 a). The data suggests little difference in crystallinity between the composites and neat PCL, or between the composites themselves. Therefore, chain shortening due to melt-processing is considered the most likely reason for the deviation from literature at NP loadings above 2 wt%. The longer compounding times used for higher NP loadings likely induced chain shortening of the PCL, a side effect of melt processing thermoplastics which can be caused by both excessive temperatures and by shearing forces [29]. Decreases in chain length of PCL have been experimentally correlated with a decreasing elastic modulus [33]. Random chain scission of PCL due to thermal energy alone is negligible below 300°C [48], so in this case chain shortening is likely a direct result of the higher processing times used for higher NP loadings. To minimize PCL chain shortening in future filament fabrication, maximum allowable processing times and auger shear forces must be identified.

### 2.4 Radiopaque Composite Biocompatibility

The biocompatibility of PCL is well established [11] and composite agents are frequently added to improve cell adhesion and growth [1, 10, 34]. However, incorporating Bi_2_O_3_ into PCL may result in prolonged cellular exposure above biocompatible levels as the composite degrades *in vivo*. To assess the impact of the range of Bi_2_O_3_ loadings used in this study on the effectiveness of TES, both initial and long-term impacts must be considered.

The initial impacts of Bi_2_O_3_ composition on cell attachment, growth, and metabolic activity of fibroblasts were assessed with a 7-day study (Fig 4). When fibroblasts were seeded onto 3D-printed scaffolds (0-11.7 wt%) for 7 days, cells proliferated on all substrates, Fig 4(a). Initial cellular attachment was lowest at 0wt% Bi_2_O_3_ (16.8 ± 13%) and highest at 0.8 wt% (54.1 ±23%). The high deviations in initial cell seeding were likely due to difficulties seeding cells onto the highly porous 3D constructs. No significant changes in cell number, as a function of NP composition, were noted on day 7. Neat PCL (0 wt%) showed the greatest overall proliferation, likely due to a low initial cell seeding. Metabolic activity had similar results, with a significant increase from day 1 to day 7 as cells proliferated. While the 2 and 6 wt% composites showed decreased metabolic activity compared to the neat PCL at early time points, there was no significant trend of decreased metabolism with increased Bi_2_O_3_ wt%. Overall, these results suggest little to no impact on early cellular proliferation, up to the maximum tested Bi_2_O_3_ loading of 11.7 wt%.

**Figure 4:**
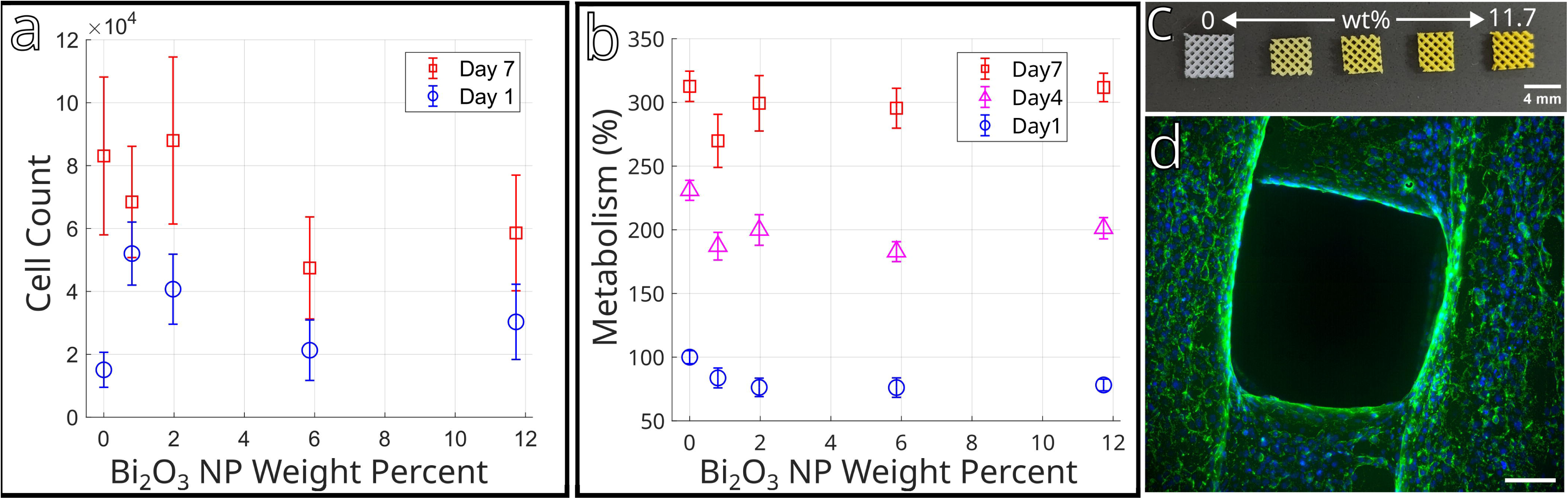
Composites of Bi_2_O_3_ NP and PCL showed excellent biocompatibility over 7 days in culture. 3T3 mouse fibroblasts seeded on 3D printed scaffolds with 0-11.7 wt% Bi_2_O_3_ had **(a)** increases in cell count for all Bi_2_O_3_ concentrations tested. **(b)** Cell metabolism increased significantly from day 1 to 7 over all Bi_2_O_3_ wt%; data was normalized to 0wt% at day 1. **(c)** 3D printed scaffolds used in cytocompatibility tests (4 × 4 mm). **(d)** Representative cell dispersion on scaffold (11.7 wt%) after 7 days, (green: actin, blue: nucleus). Data is presented as mean ± standard error. Scale bar: (c) 4 mm, (d) 100 µm.

To extrapolate the potential long-term impact of uncoated Bi_2_O_3_ NPs on cell growth and metabolism as the devices degrade, the degradation kinetics of PCL must be considered. PCL, once implanted *in vivo*, undergoes ∼2 years of hydrolytic surface and bulk degradation, during which the molecular weight decreases linearly with time [38], followed by a second phase in which phagosomes uptake the remaining low molecular weight, highly crystalline fragments [39]. The local dose of Bi_2_O_3_ observed at the site of implantation is dependent on initial Bi_2_O_3_ concentration. The formulated composites have bulk Bi_2_O_3_ concentrations ranging from 9 to 147 mg/ml (0.8 to 11.7 wt%). Assuming the NPs are exuded over the duration of device degradation, the average daily dose per unit volume (D) is roughly estimated as:

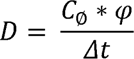

Where C_ø_ is the initial NP concentration of the composite, ((J is the as-printed volume fraction of infill vs void space, and Δt is the degradation time. For scaffolds printed with 40% infill and a conservatively estimated degradation time of 2 years, the average daily Bi_2_O_3_ NP excretion *in vivo* from the 3D printed composite scaffolds would range from 5 to 80 µg/ml per day (0.8 to 11.7 wt%).

Several recent studies have investigated the acceptable limits of Bi_2_O_3_ NP dose *in vitro*. A 2017 study investigated the effects of Bi_2_O_3_ NPs of similar size (149 nm) on mammalian liver, kidney, intestine, and lung cells. The NPs were found to increase oxidative stress, DNA damage, and cell apoptosis, with IC_50_ values ranging from 35-97*µ*g/ml depending on cell type [18]. Taking the lowest IC_50_ as the maximum allowable daily dose (35 *µ*g/ml) corresponds to a composite Bi_2_O_3_ concentration of ∼ 5.5 wt%. For more densely printed structures, the composite concentration must be further restricted to achieve an equivalent dose. However, these cell types are not likely to come into direct contact with the degrading structure. The actual concentrations observed by the liver and kidney would correspond to the overall size of the TES in addition to the bulk NP concentration, meaning smaller and less heavily loaded TES pose less risk to liver and kidney cells.

A second study in 2022 found that smaller Bi_2_O_3_ NPs (6 ± 1 nm) were effectively non-toxic to fibroblasts and murine macrophage-like cells *in vitro* with an IC_50_ of *>* 100 *µ*g/ml for both cell types [19]. This suggests the phagosome uptake of crystalline fragments could contain higher than average Bi_2_O_3_ NP concentrations without negatively affecting cell growth during the later stages of degradation. To ensure biocompatibility in clinical applications, future printing of Bi_2_O_3_ composites will focus on incorporating NPs of this size.

Overall, these results support advancing the composite as a material for TES. Additional study is required to understand the IC50 of Bi_2_O_3_ NPs for cell types relevant to orthopedic application such as osteoblasts, as well as the biological path of the NPs during and after degradation *in vivo*. Such studies would improve the estimation of the upper allowable limit of Bi_2_O_3_ in 3D printed TES.

### 2.5 X-ray Attenuation of Radiopaque Composites

A third condition for determining the optimum Bi_2_O_3_ range of incorporation was the ability to monitor composites. To that end, the relationship between Bi_2_O_3_ concentration and radiopacity was characterized by quantifying the x-ray intensity of the filaments when imaged via *µ*CT in an aqueous environment mimicking tissue. Increasing Bi_2_O_3_ was found to be directly proportional to signal intensity at a rate of 269 HU per wt% (Fig. 5 a). This is a higher rate than Arnold et al., who reported increasing radiopacity of 3D printed structures at a rate of 196 HU per wt% Bi_2_O_3_ [12]. However, Arnold et al. used a scan energy of 120 keV, compared to the 90 keV energy used here, which is the standard for preclinical *in vivo* CT. Both the lower scan energy and the proximity of the peak scan energy to Bismuth’s k-edge can explain the higher attenuation rate reported in this study. As x-ray scan energy increases, a material’s x-ray attenuation generally decreases, except around the material specific attenuation peak known as the k-edge.[20] In this study, the peak scan energy of 90 kV was very close to the k-edge of bismuth (91 keV).

**Figure 5:**
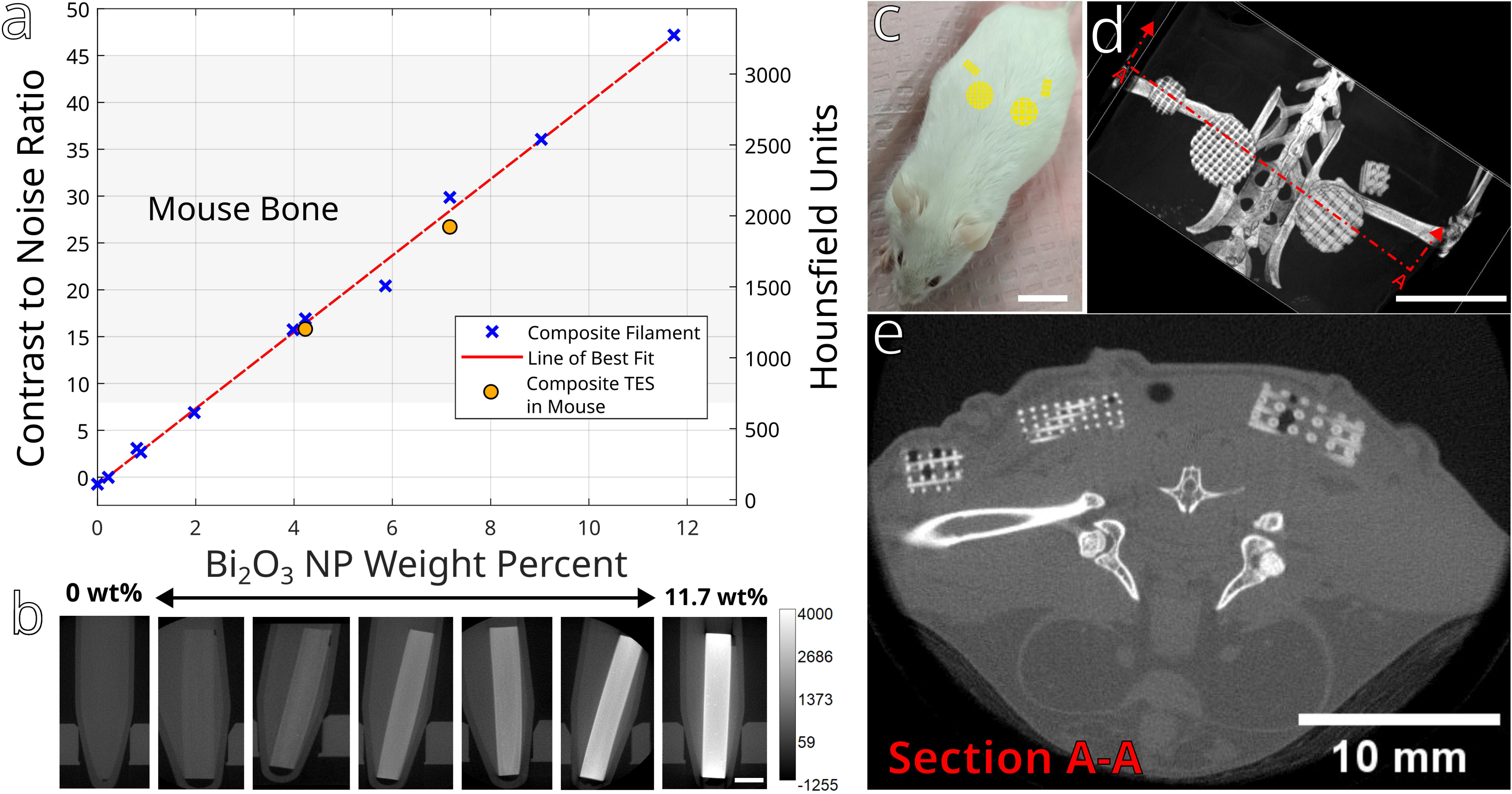
Filament radiopacity was measured via *µ*CT in both an *in vitro* aqueous environment and an *ex vivo* mouse model. Both modalities confirmed the excellent radiographic performance of the composite as low as 2 wt% Bi_2_O_3_. Increasing NP concentration correlates linearly with radiographic distinguishability shown by **(a)** plotting CNR of composites and **(b)** corresponding CT images; 4 and 7 wt% Bi_2_O_3_ TES show up like bone. **(c)** *Ex vivo* mouse study showing the approximate location of TES implantation (yellow). **(d)** 3D reconstruction from µCT of TES in mouse model. **(e)** 2D slice of the *µ*CT scan showing soft tissue, bone, and the TES. Scale bar: (b) 3mm and (c-e) 10 mm.

To verify that the attenuation calculated *in vitro* would correlate to *in vivo* studies, TES with 4.2 and 7 wt% Bi_2_O_3_ NPs were implanted into a mouse model *ex vivo* and imaged, as before (Fig. 5 c-e). The TES were easily distinguished from surrounding muscle and soft tissue, falling within the same signal range as mouse bone. To enable comparison between *in* vitro and *ex vivo* scans, the signal from the radiopaque composites was converted to a contrast to noise ratio (**CNR**). Converting the raw signal to CNR is useful as it considers the signal from the surrounding tissue in addition to the signal of the TES. A device which is indistinguishable against bone can be highly distinguishable against muscle tissue.

Comparison of the linear fitted curve of filament signal vs. wt% showed excellent agreement between the CNR of the filament and the implanted TES (Fig. 5 a). The plot shows that an equivalent CNR to mouse bone (≥ 8) could be achieved at just 2.2 wt% Bi_2_O_3_ and suggests distinguishable devices may be produced with concentrations approaching 1 wt%. This is multiple times less than the minimum weight of TaO_x_ NPs (5wt%) necessary to enable *in situ* monitoring of similar polymeric devices.[4] Similarly, it is lower than the approximately 3 wt% BaSO_4_ used in the commercially available, implantable, non-biodegradable birth control device Nexplanon [35]. However, CNR of radiopaque TES in CT has been shown to decrease when feature size falls below 5mm [21]. Therefore, imparting a sufficiently high CNR is important not only to increase initial visibility, but also to ensure long term visibility as a device undergoes degradation. Consideration of how long a device needs to be monitored should therefore impact selection of Bi_2_O_3_ wt%, especially if a range of < 2 wt% is being considered.

### 2.6 Clinical Potential of Radiopaque TES

Characterization of the 3D printed PCL-Bi_2_O_3_ NP composite produced an optimal range of 1-5.5 wt% bismuth to satisfy biocompatibility and imaging requirements. The excellent radiographic performance of the composites encouraged their demonstration at a clinically relevant scale. In orthopedics, 3D printing has been extensively studied to create anatomically sized TES post trauma [42]. An emerging application for 3D printing is for manufacturing replacement a meniscus, a tissue that acts to stabilize the knee during motion and that has a high incidence of injury during sports [22]. Patient specific devices have already been produced as alternatives to commercially available implants [22, 23, 24]. Building on these studies, an *ex vivo* model of a swine meniscus was selected for demonstration of clinical potential for radiopaque TES, due to the similarity in scale with the human meniscus [25, 26, 27]. To fabricate menisci TES, post mortem swine legs were scanned in clinical CT. The medial menisci were digitally extracted and converted to “patient specific” STL models. A model of one such swine leg and corresponding meniscus reconstruction is shown in Fig. 6 a-b. Then, the menisci were surgically extracted to prepare the pig legs for implantation of 3D printed menisci.

**Figure 6:**
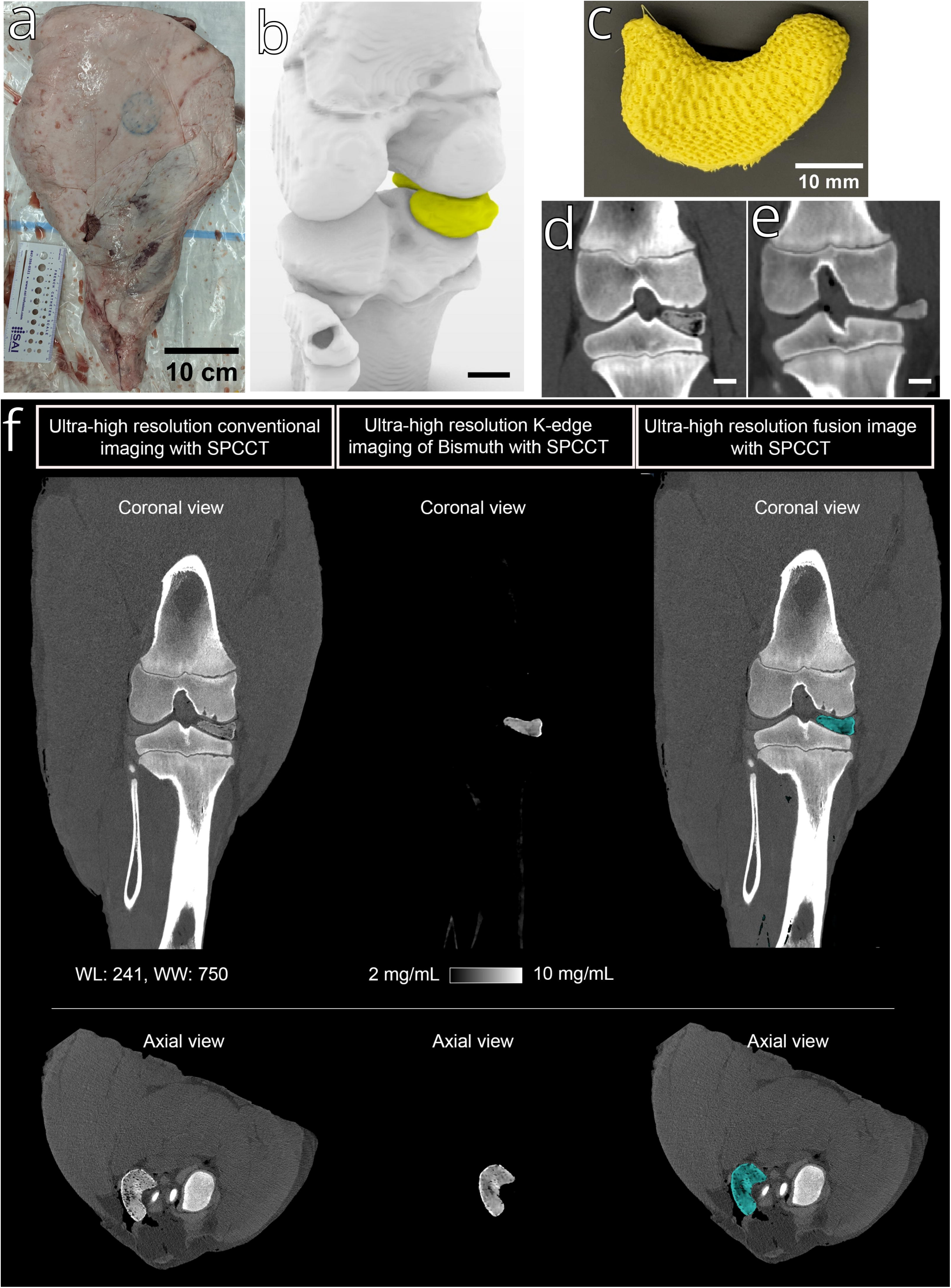
The potential to utilize radiopaque TES (PCL + 4 wt% Bi_2_O_3_) for personalized medicine was demonstrated in an ex vivo pig model. A patient specific meniscus was generated from clinical CT scans and used to 3D print a radiopaque PCL replacement. **(a)** Swine leg used in meniscal replacement. **(b)** Reconstruction of stifle joint based on clinical CT scan, medial meniscus shown in yellow. 3D printed meniscus **(c)** with 4 wt% Bi_2_O_3_ based on clinical CT scan data. Aligned **(d)** and mis-aligned **(e)** menisci imaged via clinical CT post implantation. **(f)** PCCT showing meniscus *in situ* (left), and sectioned meniscus using the unique signal resulting from the high k-edge of bismuth (91 keV) (center, right). Scale bar: (a) 10 cm, (b-e) 10 mm

The digitally extracted menisci were sufficiently large that 3D printing could be performed at room temperature ambient conditions. Due to their curved nature, the menisci could not be oriented on the 3D printer build plate to avoid overhanging features, so the “Unsupported Overhangs” parameters from Table 1 were implemented. For the uppermost layers of the menisci the “Small Cross Sections” parameters were implemented. The 4 wt% composite filament was selected to print the menisci to ensure excellent distinguishability from muscle (CNR=16).

Two menisci were 3D printed and re-implanted into the legs they were derived from. One was implanted with excellent alignment, and one was implanted in a mis-aligned orientation. The facile identification of the composite TES with clinical CT and corresponding alignment/mis-alignment were thus demonstrated and are shown in Fig. 6 d-e. In the clinic, an adverse event to meniscal retears or implant failures is difficult to diagnose. Utilizing MRI for diagnosis is difficult and generally inconclusive due to high-intensity T1 and T2 signal changes from scar tissue and greater vascularization, necessitating invasive surgical techniques instead [43, 44]. Here we demonstrate how non-invasive CT of radiopaque implants can be used for facile diagnosis.

After validation of CT monitoring to distinguish implant alignment, the ex vivo model was imaged using clinical PCCT, Fig. 6 e. Unlike traditional CT, PCCT measures the energy of transmitted x-rays and” bins” them to perform a re-convolution of the elemental distribution based on the elements k-edge. Bismuth has a distinct k-edge that is within clinical CT energy levels (91 keV) [8]. The composite medial meniscus implant is readily identifiable in the ultra-high-resolution scan, Fig. 6 f. The voxels above the Bi_2_O_3_ threshold content were isolated to generate the reconstruction of the anatomically derived, 3D printed radiopaque meniscus. The fusion image demonstrates excellent radiographic distinguishability, even in applications where implants possess similar x-ray attenuation to bone in a conventional CT. With the growth of clinical PCCT, the incorporation of metal oxide-based NPs is a technology poised to bring enhanced diagnostic power and longitudinal traceability to implanted biomedical devices.

## 3 Conclusion

The movement towards personalized on-demand biomedical implants manufactured at the point-of-care is a key driver for 3D printing of biomedical devices. Ideally, devices would also incorporate the ability to be monitored in situ by clinical imaging modalities, such as CT. We demonstrate a significant step towards this goal by fabricating and 3D printing radiopaque PCL filament incorporating Bi_2_O_3_ NPs. In the process, we also present an optimized method for FFF 3D printing of PCL TES accommodating the low recrystallization temperature of PCL and composites. All NP composites had good mechanical properties and biocompatibility. Imaging via *µ*CT showed an increase of 4.1 in CNR with each additional wt.% Bi_2_O_3_ incorporated. Above 2.2 wt.% Bi_2_O_3_, composites were radiographically distinguishable from soft tissue, with CNRs in the range of mouse bone. Further, the clinical potential of radiopaque 3D printed TES was demonstrated in an ex vivo orthopedic application. Replacement swine menisci were 3D printed from radiopaque composite with anatomical accuracy and once implanted, enabled facile identification of device alignment at a low NP concentration (4 wt% Bi_2_O_3_). Not only can Bi_2_O_3_ composite TES be distinguished using traditional CT, but they are also well suited to the emerging clinical imaging modality of PCCT. Once imaged via PCCT, the radiopaque TES were segmented from biological structures based on the unique x-ray attenuation properties of bismuth. Together these results show the excellent performance of Bi_2_O_3_ NPs as a radiopaque additive at low concentrations (1 to 5.5 wt%) in 3D printed composite TES for longitudinal monitoring of pre-clinical and clinical devices.

## 4 Methods

### 4.1 Materials

Unless noted, materials were purchased from Fisher Scientific.

Bi_2_O_3_ NPs (99% purity) were purchased from American Elements. Polycaprolactone Filament (Facilan ™PCL 100), was purchased from 3D4Makers (50kg/mol, 2.85±0.1 mm). Dulbecco’s modified Eagle’s medium (DMEM, 11965), fetal bovine serum (FBS, 160,000,036), DAPI Stain (62248), and Penicillin–Streptomycin (15-140-122) were purchased from Thermofisher Scientific. AlamarBlue (cat 786-921) was purchased from G-Biosciences. ActinRed 555 ReadyProbes reagent (R37112) and Slow Fade Diamond Antifade Mountant (S36972) were purchased from Invitrogen. Papain Buffer (0.1M phosphate buffer, 10mM L-cysteine, and 2mM Ethylenediaminetetraacetic acid (EDTA), 3U/ml papain) were purchased from Sigma-Aldrich.

### 4.2 Radiopaque Filament Production

Bi O NPs were combined with Facilan™PCL 100 filament in dichloromethane at a ratio of 50 wt% NP to PCL. As a result of adding yellow Bi_2_O_3_ NPs, the fabricated filaments were easily distinguishable from the purchased PCL filament by color. The solution was vortexed and then sonicated for 5 minutes before being cast onto a glass plate and dried overnight in a fume hood. The cast and dried 50 wt% precursor was cut into small squares (*<* 5mm) and compounded with additional PCL in a DSM 15 CC dual auger micro compounder to nominal concentrations of 1.5 to 20 wt% Bi_2_O_3_. The dual auger micro compounder was run at a rate of 100 RPM. During compounding, each composite was recycled for at least 10 minutes at 100 RPM and 165°C under argon gas to ensure homogenous NP dispersion. %. To ensure NP dispersion at higher loadings, composites with the highest nominal concentrations were compounded for 40 min at 165° C and compounding times decreased with nominal NP content down to 10 minutes. The resulting composites were extruded into filament using a Filabot EX2 single auger extruder. The filament was first extruded at 120°C into a water bath, dried, chopped, and then recycled at least twice to ensure homogeneity. Finally,, dimensionally accurate filament was extruded under the same conditions at 2.85 ± 0.1 mm. The process can be seen graphically in 1(a).

### 4.3 Inductively Coupled Plasma Mass Spectrometry (ICP-MS)

ICP-MS was performed in the MSU Quantitative Bio-Element Analysis and Mapping Center (QBEAM) on an Agilent 8900. First, 10mg samples of each filament were dissolved in 10 ml of 70% HNO_3_. Next, the ICP-MS was run with a wash and calibrated with known quantities of Bi-209 against an In-115 standard. A wash was run again before running the samples of dissolved filament. To calculate the Bi_2_O_3_ wt% of each filament, the volume of each solvate was first calculated using the equation

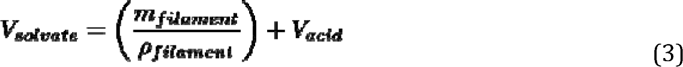

where *V* is volume, *m* is mass, and *ρ* is density. From the ICP-MS results, the final Bi_2_O_3_ weight percent was calculated as

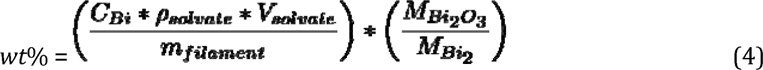

where *C* is the bismuth concentration and *M* is the molar mass.

### 4.4 Differential Scanning Calorimetry (DSC)

DSC was performed on a TA Instruments DSC2000. 10mg samples were analyzed in aluminum sample pans. Samples were heated and cooled at a rate of 10 ^⍰^C/min. The samples were heated to 100^⍰^C in order to eliminate any processing effects. Data logging then began and samples were cooled down to -10^⍰^C, then heated back up to 100^⍰^C. The data was plotted in MATLAB and the maxima and minima were reported as the recrystallization and melting temperatures, respectively.

### 4.5 Scanning Electron Microscopy (SEM)

SEM was performed on a TESCAN MIRA3. Samples of composite filament were sputter-coated with Pt. Imaging was performed with backscattered electrons at an accelerating voltage of 20 keV.

### 4.6 Transmission Electron Microscopy (TEM)

TEM was performed on a JOEL 1400 at 100 keV at the MSU Center for Advanced Microscopy. Bi_2_O_3_ NPs were dispersed in ethanol then deposited on a mesh grid and dried in air before being inserted into the TEM for imaging.

### 4.7 3D Printing

All 3D printing was performed on a Lulzbot Taz 6 using 2.85mm diameter filament with either a 0.5 or 0.3 mm nozzle. Modified cooling fan shrouds were designed and installed to direct flow closer to the immediate area of extrusion. All TES were printed with rectilinear infill patterns at 40-50% infill. Optimized printing parameters can be seen in Table 1. All STL models were prepared for 3D printing using PrusaSlicer 2.6.1 opensource software.

### 4.8 Tensile Testing

Tensile testing was performed on a Stable Micro Systems TA-XT plus with a 5kg load cell. 3mm wide tensile specimens were cut from 3D printed mats of dimensions 1mm deep x 25mm long x 50 mm wide (40% infill, rectilinear infill). The mats were 3D printed according to “Cold Room” parameters as they were sufficiently small to achieve adequate bed adhesion. The solid edges of the tensile samples were clamped at both ends using textured tensile grips. The specimens were tested to failure, n=5. The data is represented as an average stress-strain curve for each composite in Fig. 3 a. Yield stress and strain are represented as mean ± standard deviation.

### 4.9 Biocompatibility

3T3 embryonic mouse fibroblasts (ATCC) were cultured in complete media (DMEM, 10% FBS, 1% Penicillin– Streptomycin) and incubated at 37°C, 5% CO_2_ for 7 days. The scaffold samples (4 × 4 mm) were cut from 3D printed sheets (40% infill, 1 mm thick). The samples were sterilized in 70% ethanol for 1 hour then washed in sterile water. After immersion in complete media in a 48 well plate, the samples were incubated for 1 hour and moved to empty wells for seeding. 1 x 10^5^ cells in 5*µ*l complete media were pipetted onto each scaffold and the scaffolds were incubated for 3 hours, then moved to empty wells with 300*µ*l complete media. Additionally, a cell standard 0 to 5 × 10^5^ cells was collected and frozen at -80^⍰^C for later comparison.

Metabolic activity was tested at days 1, 4 and 7. Each scaffold was incubated in complete media with 10% AlamarBlue for two hours. The media was removed and its fluorescence was read with a BioTek plate reader (Ex = 530 nm, Em = 590 nm). After incubation with AlamarBlue, the scaffolds were replaced in complete media for continued culture until day 7.

Cell count was measured on days 1 and 7 using a using Quant-iT™PicoGreen™dsDNA Assay Kit (ThermoFisher Scientific). The scaffolds were washed for 5 minutes in PBS and then frozen at -80^⍰^C. After thawing, the samples were digested overnight in 200*µ*l of papain buffer at 60^⍰^C, then diluted 1:4 in TNE buffer before loading into a black 96 well plate. PicoGreen dye (1:200 in TNE buffer) was added and the fluorescence was read (Ex = 480 nm, Em = 520 nm). The DNA content for each scaffold was determined by comparison to the cell standard collected on day 0 and a DNA standard provided by the kit (1 *µ*g/ml - 1 mg/ml).

All tests were repeated twice, totaling n=10 for metabolism and n=6 for cell count, and presented as mean ± standard error.

### 4.10 Immunohistochemistry

To visualize the distribution of 3T3 cells on the 3D printed scaffolds, samples were imaged via light microscopy. To prepare the samples, 3T3 cells were seeded onto scaffolds, as before. On day 7, they were fixed for 15 minutes in 4% paraformaldehyde and stored in PBS at 4^⍰^C. Samples were washed in PBS+0.1% Tween-20 (PBST) and permeabilized in 0.1% Triton X-100 in PBS, followed by two PBST washes. Blocking was done with 5 wt% BSA in PBST (1 hr, 23^⍰^C) with two PBST washes. The actin cytoskeleton was stained with ActinRed 555 per manufacturer’s instructions. Samples were washed twice with PBS and left in a solution of PBS and DAPI stain.

Stained films were inverted onto a coverslip in a drop of Slow Fade Diamond Antifade Mountant. Imaging was performed using a Leica DMi8 microscope with LAS X software. Z-stacks were taken and post-processed with Thunder imaging system (Leica) to remove background fluorescence.

### 4.11 X-Ray Computed Tomography (Micro-CT)

The *µ*CT scans were performed on a Perkin Elmer Quantum GX at 90 kV, 88*µ*A. DICOM files were processed with the FIJI distribution of Image J. Contrast to noise (CNR) was calculated as

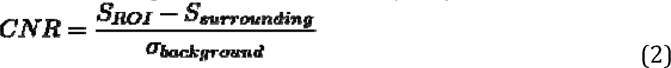

where S are signal intensities of the region of interest (ROI) and surrounding tissue in Hounsfield units (HU), and *σ* is the standard deviation (noise) of the scan in HU.

Clinical CT scans were performed on a GE 64-Slice CT utilizing the pre-set clinical parameters for orthopedics. DICOM images were processed in Image J and ITK Snap.

### 4.12 Meniscus Segmentation

The menisci of the pig legs were segmented out from the DICOM files collected via clinical CT using the active contour segmentation tool in ITK Snap software. First, the signal threshold was carefully set to isolate the signal from the meniscus from the surrounding bone and tissue (40 *<* Signal *<* 122 HU). Next, areas of each meniscus were initialized with seed regions. Third, the seed regions were grown iteratively to segment the entire meniscus. Finally, the segmented regions were exported as STL files. The same process was used to segment out the bones surrounding the meniscus. The segmented menisci were reconstructed with the menisci to confirm proper segmentation and anatomical fit.

### 4.13 Surgical Implantation

The medial meniscus was surgically removed from the corresponding stifle joint of a pig leg, and the 3D printed meniscus was anchored in place with surgical glue with surrounding tissues sutured closed. Implants were secured both in and out of alignment.

### 4.14 Photon Counting Computed Tomography (SPCCT)

A clinical photon-counting CT (PCCT) system prototype (Philips, Haifa, Israel) was used to scan the samples. This system is equipped with a 2 mm thick CZT, bonded to Philips’ proprietary ChromAIX2 application-specific integrated circuit [30]. The system enables a 500 mm in-plane field-of-view, and a z-coverage of 17.6 mm in the isocenter, realized with 64 rows of detectors, 270×270 µm² in size. The application-specific integrated circuit (ASIC) supports the readout of 5 different energy bins, which were at 30, 51, 63, 78, and 92 keV for the scans of the bismuth contrast agents in order to differentiate its attenuation profile and to coincide with its K-edge at 90.5 keV [31]. More technical details of the reconstruction chain are provided in a previous study [6].

Acquisition parameters were similar to a clinical protocol, using a tube voltage at 120 kVp, a tube current at 182 mA, exposure of 190 mAs and a computed tomography dose index at 19.2 mGy. Reconstruction parameters were set to provide both conventional HU images and K-edge images of the bismuth, with a field of view of 256 mm, a matrix size of 1024 × 1024 pixels an isotropic voxel size of 250 μm and using a detailed bone (sharp) reconstruction kernel (e.g., Detailed 2).

### 4.15 Statistics

One-way ANOVA statistics were performed on tensile testing in Matlab. One and Two-way ANOVA statistics were performed on cell study data in Matlab. Results were considered statistically significant when p<0.05, and highly statistically significant when p<0.001.

## Acknowledgements

The authors would like to thank Sabrina Johnson, Ed Drown, Bryce Borgialli, and Legend Kenney. This study was supported by the National Institute of Biomedical Imaging and Bioengineering of the NIH under award number R01 EB029418. The content is solely the responsibility of the authors and does not necessarily represent the official views of the National Institutes of Health.

